# Brightness illusions evoke pupil constriction and a preceding primary visual cortex response in rats

**DOI:** 10.1101/2022.07.13.499566

**Authors:** Dmitrii Vasilev, Isabel Raposo, Nelson K. Totah

## Abstract

The mind affects the body via central nervous system (CNS) control of the autonomic nervous system (ANS). In humans, one striking illustration of the mind-body connection is that illusions, subjectively perceived as bright, drive constriction of the eye’s pupil by activating the sympathetic arm of the ANS. How the CNS is involved in this pupil response is unknown and requires an animal model for intracerebral investigation of potential regions, cell types, and neuronal projections. However, the physiological response to this illusion has long been thought to occur only in humans. Here, we report that the same brightness illusion that evokes pupil constriction in humans also does so in rats. Cortex-wide EEG recordings revealed that, compared to a luminance-matched control stimulus, the illusion (which appears subjectively brighter to humans) evoked a larger response only in primary visual cortex (V1). This cortical response preceded pupil constriction by ~335 msec suggesting a potential causal role for V1 on the pupil. Our results establish a new animal model of importance for studying how the CNS response involved in sensing a brightness illusion drives a physiological reaction in the body. We provide objective evidence that complex mind-body connections are not confined to humans and that V1 may be part of a shared, mammalian, neural network for bodily reactions to illusions.

## Introduction

Mental processes mediated by the central nervous system (CNS) can affect the body via the autonomic nervous system (ANS). The mind-body connection is apparent from the effects of psychological stress on immune and gastrointestinal function^1–3^, and placebo effects on pain driven by a person’s beliefs^4^. One fascinating mind-body connection is that subjective illusions of brightness cause the eye’s pupil to constrict^5^ due to the CNS driving the sympathetic arm of the ANS which constricts the pupil^6^. The brain regions, cell types, and synaptic projections that mediate this mind-body interaction remain completely unexplored because there is no animal model permitting intracerebral (invasive) investigation. It has been assumed that an animal model is not possible because this type of mind-body connection occurs only in humans.

In this study, we presented a brightness illusion (which humans subjectively perceive as bright^5^) to head-fixed rats while performing pupillometry and simultaneous brain-wide 32-electrode EEG. We used the Asahi stimulus (**Fig. 1**), which was created by Prof. Akiyoshi Kitaoka (Department of Psychology, Ritsumeikan University, Osaka, Japan) based on luminance-gradient stimuli in earlier work^7^. We show that the Asahi stimulus drives a pupil constriction in rats. On the other hand, a luminance-matched control stimulus, which does not evoke illusory brightness in humans, did not cause a pupil constriction in rats. The Asahi stimulus also drove a larger cortex EEG event-related potential that was confined to primary visual cortex and preceded pupil constriction. Our results show that the rat is a viable animal model for studying how neural processing of illusions can affect autonomic control of the body.

**Figure 1.**
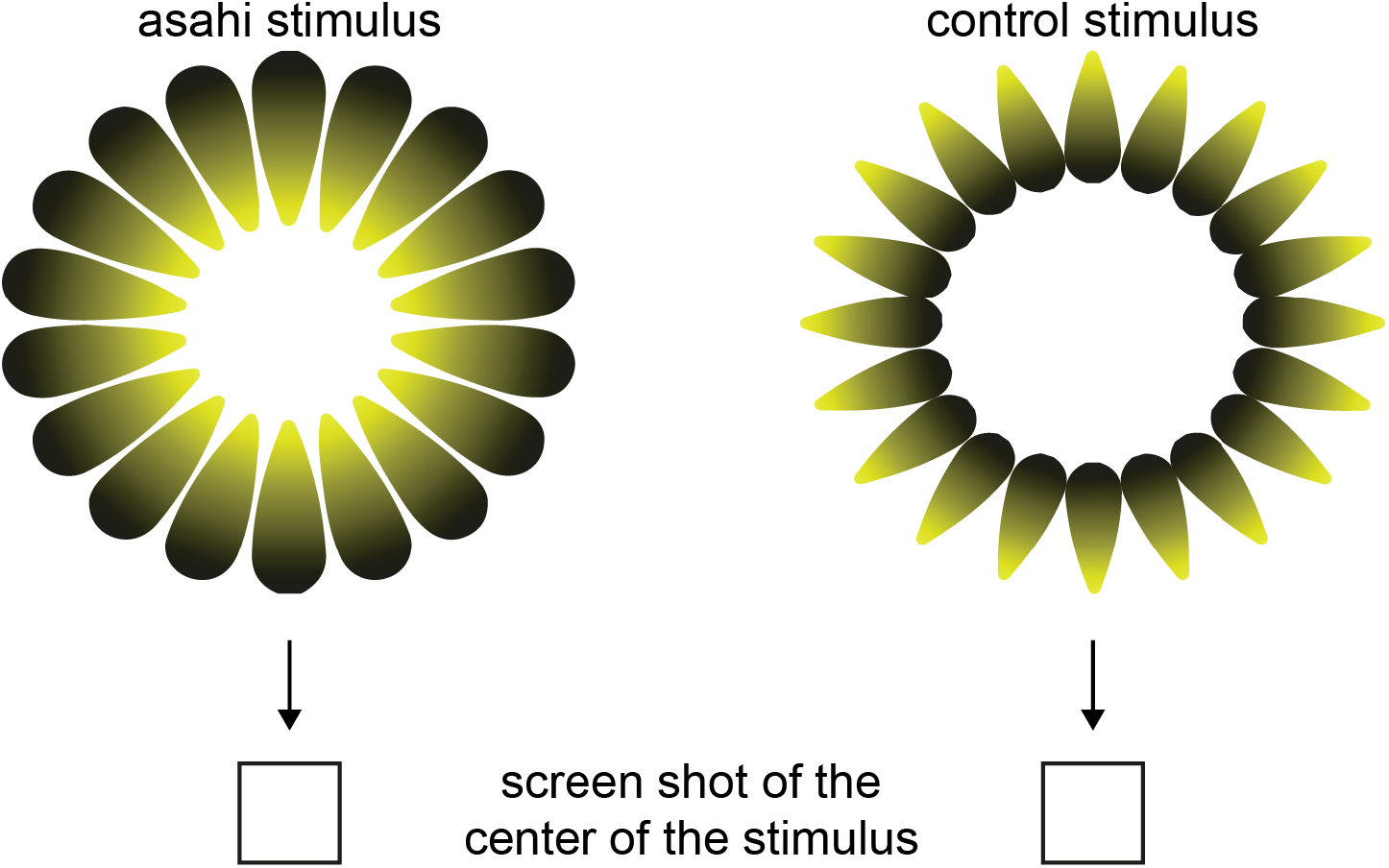
Illustrations of the Asahi stimulus and the luminance-matched control stimulus. The Asahi stimulus is typically perceived by humans to have a brighter-than-white glare in the center. By rearranging the stimulus to have a different shape, this brightness illusion is abolished. The screen shots show that the actual luminance at the center of each stimulus is identical, despite appearing to be brighter in the center of the Asahi stimulus.

## Results

Fourteen, male, lister-hooded rats were head-fixed on a non-motorized treadmill and passively exposed to visual stimuli. Rats were free to walk or sit immobile during the experiment. We presented the Asahi stimulus (**Figure 1**), which humans perceive to have a bright glare in the center^5^. We also presented a luminance-matched control stimulus (**Figure 1**) that due to rearrangement of the Asahi stimulus into a new structure, does not evoke a brightness percept in humans^5^. Although yellow stimuli (~580 nm wavelength) were used in the human study^5^, the spectral sensitivity of the rat retina is diminished around 580 nm and lacking sensitivity over 590 nm, while it is very sensitive below 530 nm^8^. Therefore, in addition to presenting the yellow stimuli from prior work in humans, we also presented 508 nm (green) stimuli (**Supplementary Figure 1**). The stimuli were various sizes (10°, 20°, 40°, and 60°) and presented by tiling the entire visual field with as many stimuli as possible. An example of the 10° yellow Asahi stimulus and luminance-matched control stimulus are shown in **Supplementary Figure 2** and **Supplementary Figure 3**, respectively.

Pupil size was measured using video frames collected at 45 frames per sec. Environmental illumination was held constant throughout the experiment and across subjects by collecting the data in closed faraday cages with total darkness except for the stimulus screen. Each stimulus was presented 50 times in randomized order for 4 sec with a 4 to 8 sec inter-stimulus interval drawn from a distribution with a flat hazard rate. Between stimuli, we presented a grey screen that was equiluminant with the stimuli (15 lux measured at the head of the rat). The inter-stimulus intervals allowed ample time for the pupil to return to baseline and the flat hazard rate reduced expectation of stimulus onset.

### The Asahi illusion evokes a pupil constriction in rats

Given the spectral sensitivity of the rat retina, we tested the hypothesis that the green Asahi illusion, but not the yellow version, would cause a pupil constriction in rats relative to the luminance-matched control stimulus. Since stimulus onset time was difficult to predict, we expected to observe a dilation of the pupil^9–11^. However, this dilation may be counteracted by competitive drive from parasympathetic activation of the sphincter pupillae muscle.

We observed a constriction of the pupil after onset of the 10° and 20°, green Asahi stimuli (**Figure 2**). The magnitude of constriction was larger for the Asahi stimulus compared to the control stimulus in those stimulus conditions. A Bayesian one-tailed t-test of the hypothesis that the magnitude of constriction was larger for the Asahi stimulus was strongly supported by these data (green, 20° stimuli: BF10 = 14.68; green, 10° stimuli: BF10 = 6.36). The SEM of the constriction latency was 578.6±67.6 msec for the green 10° Asahi stimulus and 575.4±61.8 msec for the green 20° Asahi stimulus.

**Figure 2.**
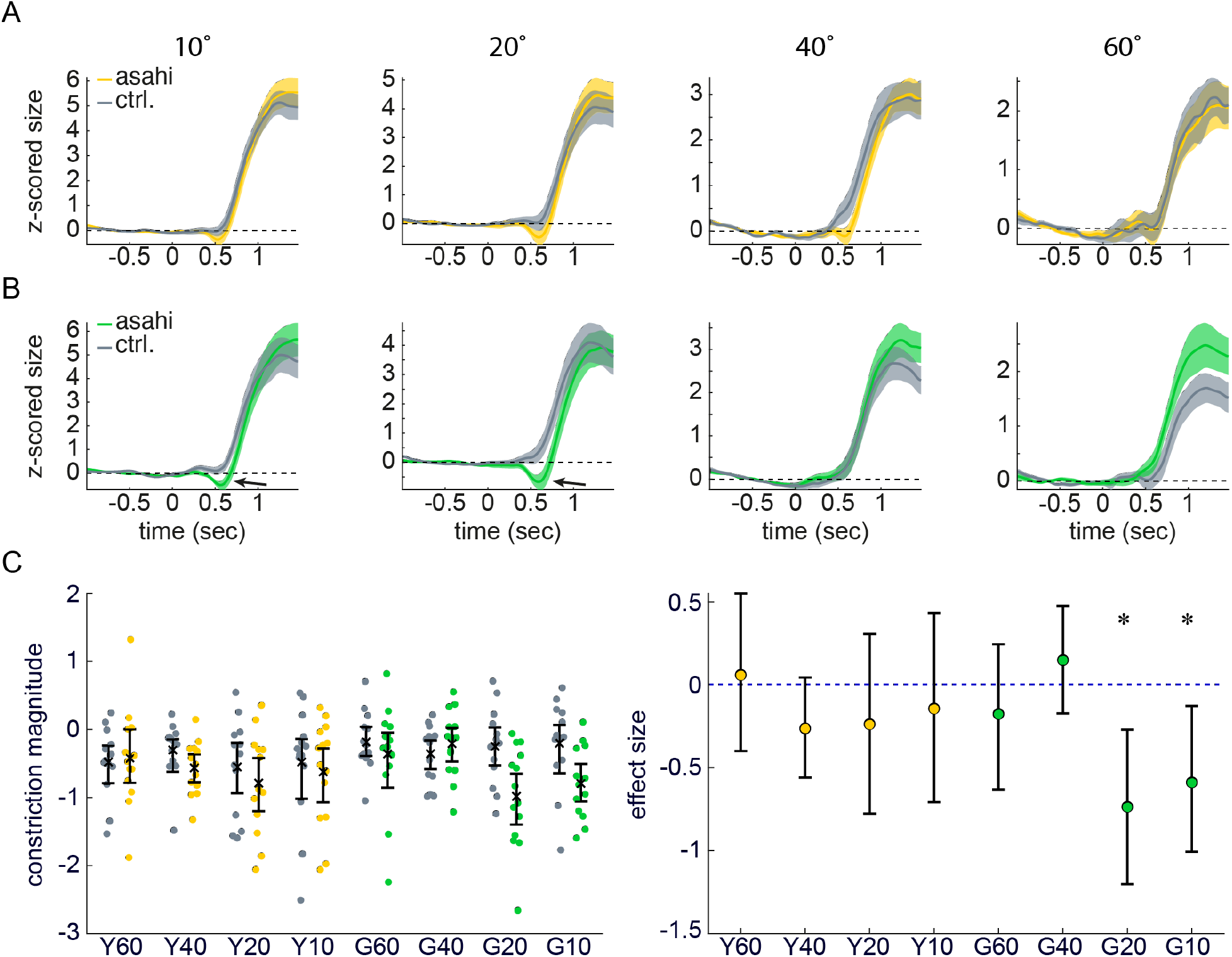
The Asahi stimulus evokes a pupil constriction in rats. **(A, B)** The SEM of pupil size is plotted from 1 sec before stimulus onset until stimulus offset at 4 sec later. Pupil size was normalized relative to the pre-stimulus pupil size using a z-score. The grey line is the pupil size around the control stimulus and the colored line is relative to the Asahi stimulus. Panel A shows the pupil response to yellow stimuli and panel B shows the response to green stimuli. A constriction relative to baseline is apparent for 10° and 20° green stimuli (arrows). **(C)** The minimal pupil size (z-scored) in the first 750 msec after stimulus onset is plotted for each stimulus. Dots are individual rats. In the yellow 60°, 40°, and 20° conditions and the green 10° condition, only 13 rats are plotted due to data loss. The right panel shows the effect size between the Asahi stimulus and the control stimulus. The asterisks indicate support for the alternative hypothesis (BF10 > 3).

We consider it unlikely that fixation on local changes in contrast could explain the observed pupil constriction. Fixation on local contrast changes is improbable because rats do not have foveal vision^12^ and the visual acuity of rats is lower than sharp changes in local contrast of the 10° and 20° stimuli^13^’^14^. Moreover, the same black-to-white contrast changes occur in the yellow and green stimuli, yet only the green Asahi stimuli were associated with a pupil constriction. However, we additionally compared eye position found that they did not differ between the two stimuli during the first 600 msec after stimulus onset, which was the window during which the pupil constriction was observed. Bayesian paired t-tests comparing the location of the center of the pupil between the green Asahi stimulus and green luminance-matched control stimulus supported the null hypothesis that location did not differ (x-position for 20° and 10° stimuli were BF10 = 0.34 and BF10 = 0.34, respectively; y-position for 20° and 10° stimuli were BF10 = 0.29 and BF10 = 0.34, respectively). Moreover, saccades were rare and saccade rate did not differ between stimulus conditions. The 20° and 10° Asahi stimuli had saccade rates of 0.597±0.062 and 0.859±0.083, whereas the 20° and 10° luminance-matched control stimuli had saccade rates of 0.602±0.060 and 0.834±0.079 (SEM). The observed saccade rates strongly supported the null hypothesis that mean saccade rate was similar for the Asahi stimulus and the luminance-matched control stimulus for both the 20° stimuli or the 10° stimuli (BF10 = 0.15 and 0.12, respectively).

### The visual cortex responds to the Asahi stimulus in rats

We next assessed whether the Asahi stimulus differentially engaged any cortical regions(s) in comparison to the luminance-matched control stimulus. In 10 of the 14 rats, we recorded EEG from anterior frontal cortex to visual cortex using a 32-electrode array implanted directly onto the skull and aligned to bregma. We obtained event-related potentials (ERPs) for each electrode in a window starting 750 msec before stimulus onset and lasting until 750 msec after stimulus onset. A topographical plot revealed that the response was confined to posterior electrodes laying over visual cortex. The Asahi and the control stimuli evoked a visual cortex response for both yellow and green stimuli of all sizes (**Supplementary Figure 4A,4B**). A Bayesian one-sided t-test of the hypothesis that the ERP peak was larger for the “brighter” stimulus (i.e., the Asahi stimulus) in comparison to the control stimulus was supported in the case of the 10° green Asahi stimulus (BF10 = 5.67, **Figure 3A, 3B**). Notably, this the 10° green Asahi stimulus also evoked a pupil constriction. The SEM of the ERP peak latency after stimulus onset was 244.1 ±8.0 msec for the green 10° Asahi stimulus. A topographical plot of the ERP peak magnitude shows that it was confined, not to all visual regions, but specifically to primary visual cortex (V1) for the Asahi and the control stimuli (**Figure 3C**). These electrodes were located at 5.0 mm and 7.0 mm posterior to bregma with 3 electrodes (per hemisphere) at each posterior location situated laterally from bregma at 1.5 mm (overlaying V2MM), 3.0 mm (overlaying V2ML at the anterior electrode and V1M at the posterior electrode), and 4.4 mm (overlaying V1 at the anterior electrode and V1 B at the posterior electrode). The ERP was largest over V1.

**Figure 3.**
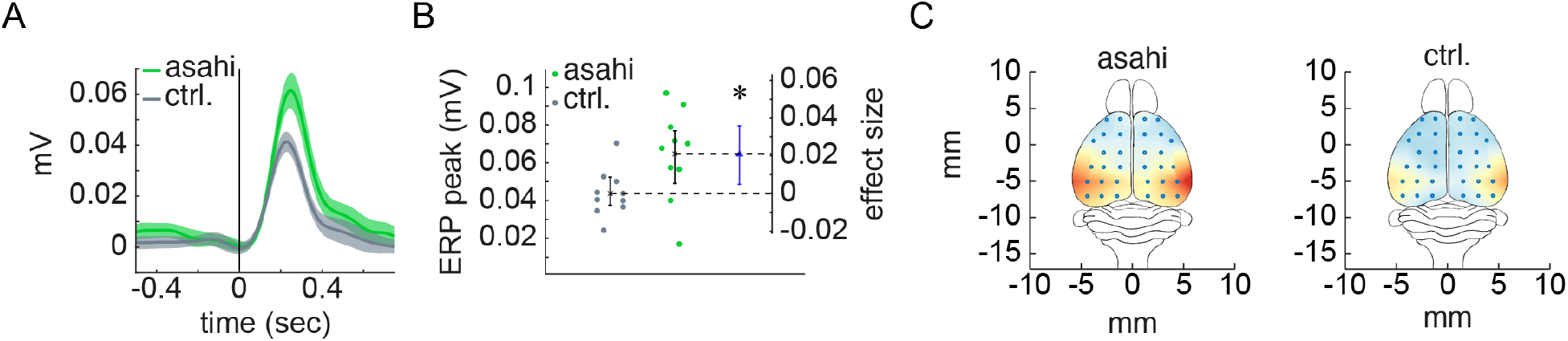
An Asahi stimulus that evoked pupil constriction also evoked a larger event-related potential in visual cortex. **(A)** The SEM of the visual cortex EEG is plotted from 750 msec before and until 750 msec after stimulus onset. The grey line is the ERP around the control stimulus and the colored line is relative to the green 10° Asahi stimulus. **(B)** The maximal potential in the ERP is plotted for each stimulus. Dots are individual rats. The inset shows the effect size between the Asahi stimulus and the control stimulus in mV. The asterisk indicates support for the alternative hypothesis (BF10 > 3). **(C)** The average ERP peak magnitude across 10 rats is shown on a scale of 0 mV (blue) to 0.06 mV (red). Electrode locations are shown relative to bregma at the origin.

## Discussion

The effect of brightness illusions on autonomic control of the pupil are a powerful tool for studying how parts of the CNS involved in higher mental functions can affect the body^5^. The neuronal correlates of this mind-body connection are unknown, and their discovery requires an animal model that permits probing cell types and specific neuronal projections using single cell recordings and optogenetic tagging of neurons by their projection target. However, it has long been assumed that this capacity for brightness illusions to evoke pupil constriction is limited to humans.

Here, using a novel combination of head-fixation, pupillometry and EEG recordings in rats, we show that the pupil constricts after the same brightness illusion that causes pupil constriction in humans. Our results establish the rat as an animal model for studying how sensing a brightness illusion drives a physiological reaction in the body. Although there was no behavioral report of perceived brightness by the rats, this paradigm allows connecting changes in the body with a high-level mental process^15^, in this case, the processing of the sensory information present in the Asahi stimulus. The processing of sensory information has been termed “sensing” and is distinct from “perception” which can be thought of as imbuing an interpretation on sensory information, or awareness of sensory information^16^. Our findings link the mental process of sensing illusory brightness to a change in body state (pupil size).

In support of this mental process interceding in the normal autonomic (and automatic) control over the pupil, which is a very fast reflex mediated by a two-synapse retina-brainstem-ANS loop^17,18^, we found that the pupil constriction to illusory brightness was strongly delayed in comparison to the pupillary light reflex. The reflexive pupil constriction to an actual increase in luminance requires approximately 250 msec after stimulus onset for high intensity light and as much as 400 msec for very low intensity light^19–21^. However, the brightness illusion (10° green Asahi stimulus) required even longer to evoke constriction (579 msec) than even very low intensity light. The delayed constriction may be due to the time (additional synapses) required for this mental process to physiologically intervene in the two-synapse ANS pupillary light reflex.

### V1 may be a key node in a shared mammalian neural network for bodily reactions to illusions

We found that V1 and, importantly no other cortical regions, responded to the brightness illusion as though it were physically brighter than the control stimulus. Visual cortex was likely to respond because illusions require the processing of the gestalt of a visual scene^22^ and such high-level visual processing relies on visual cortex in non-human primates and in mice^23–26^. Surprisingly, however, other cortical regions did not respond more strongly to the brightness illusion. For instance, although frontal regions have been implicated in the awareness and interpretation of stimuli^27^ and can activate sympathetic drive on the pupil via monosynaptic input to brainstem noradrenergic neurons^10,28–30^, frontal cortex was not particularly responsive to the brightness illusion. However, it is also possible that the activity of small populations of neurons in frontal cortex (or other cortical regions) is involved in the processing of the brightness illusion, but that their activity is not apparent in the EEG signal.

Our data are consistent with the idea that V1 may be a potential cause of pupil constriction, given that the V1 response preceded pupil constriction. The latency of the maximal ERP after the 10° green Asahi stimulus was 244 msec, whereas the pupil did not constrict until 579 msec after this stimulus. V1 may control the ANS via either an unknown direct projection or via a poly-synaptic set of subcortical synapses. These forebrain circuits must eventually modulate one or two brainstem synapses involved in this basic reflex: either the retinal ganglion cell synapses onto the midbrain olivary pretectal nucleus, or its synapses onto the preganglionic Edinger-Westphal nucleus that activates the ciliary ganglion of the parasympathetic nervous system^17,18^. However, it is imperative to test the role of V1 using optogenetic inhibition, as well as record activity in projection-target defined V1 neurons (using opto-tagging) and in potentially involved sub-cortical structures. These experiments are impossible in humans but are now permissible using this rat model.

### A new animal model of importance for studying mind-body interaction

By demonstrating that the same brightness illusion that drives pupil constriction in humans also does so in rats, we show that this type of mind-body connection is present at an earlier stage of evolution than previously thought. This finding establishes the first animal model for studying how the CNS response involved in sensing a brightness illusion drives a pupil response, which can be used to uncover the physiological basis of this mind-body interaction.

## Supplemental figures

**Supplementary Figure 1.**
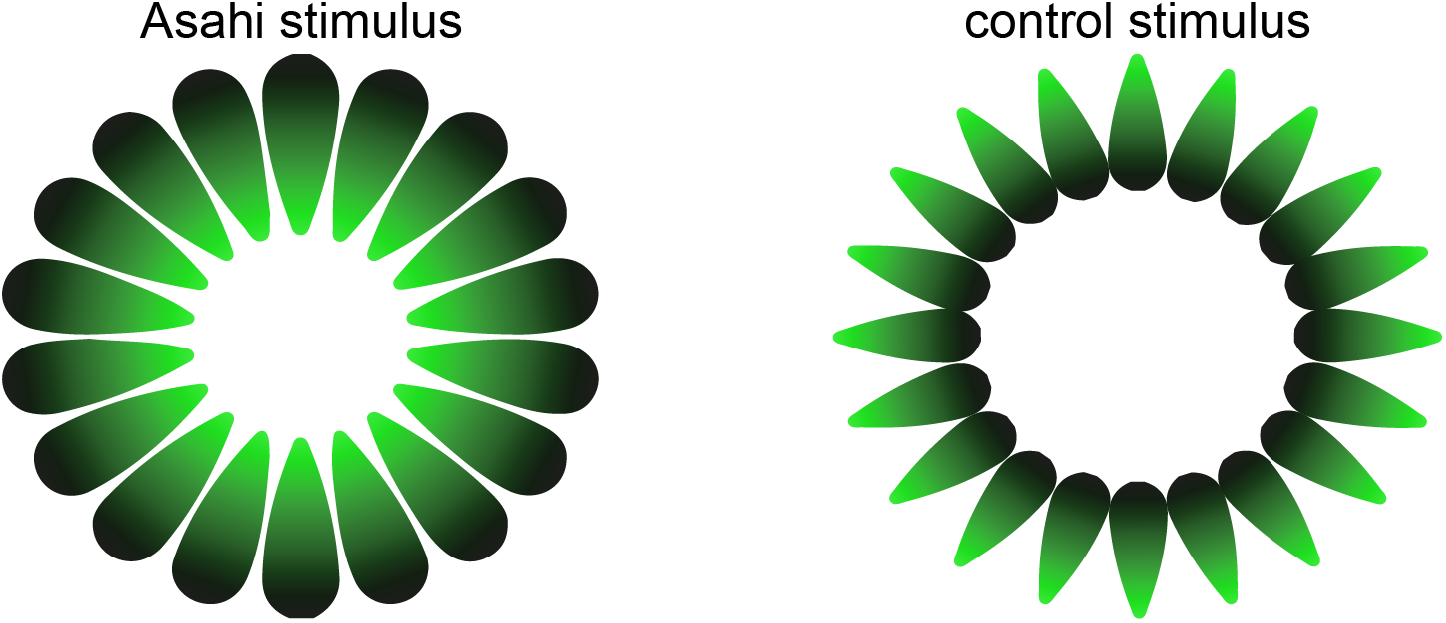
The green version of the 60° Asahi stimulus and control stimulus.

**Supplementary Figure 2.**
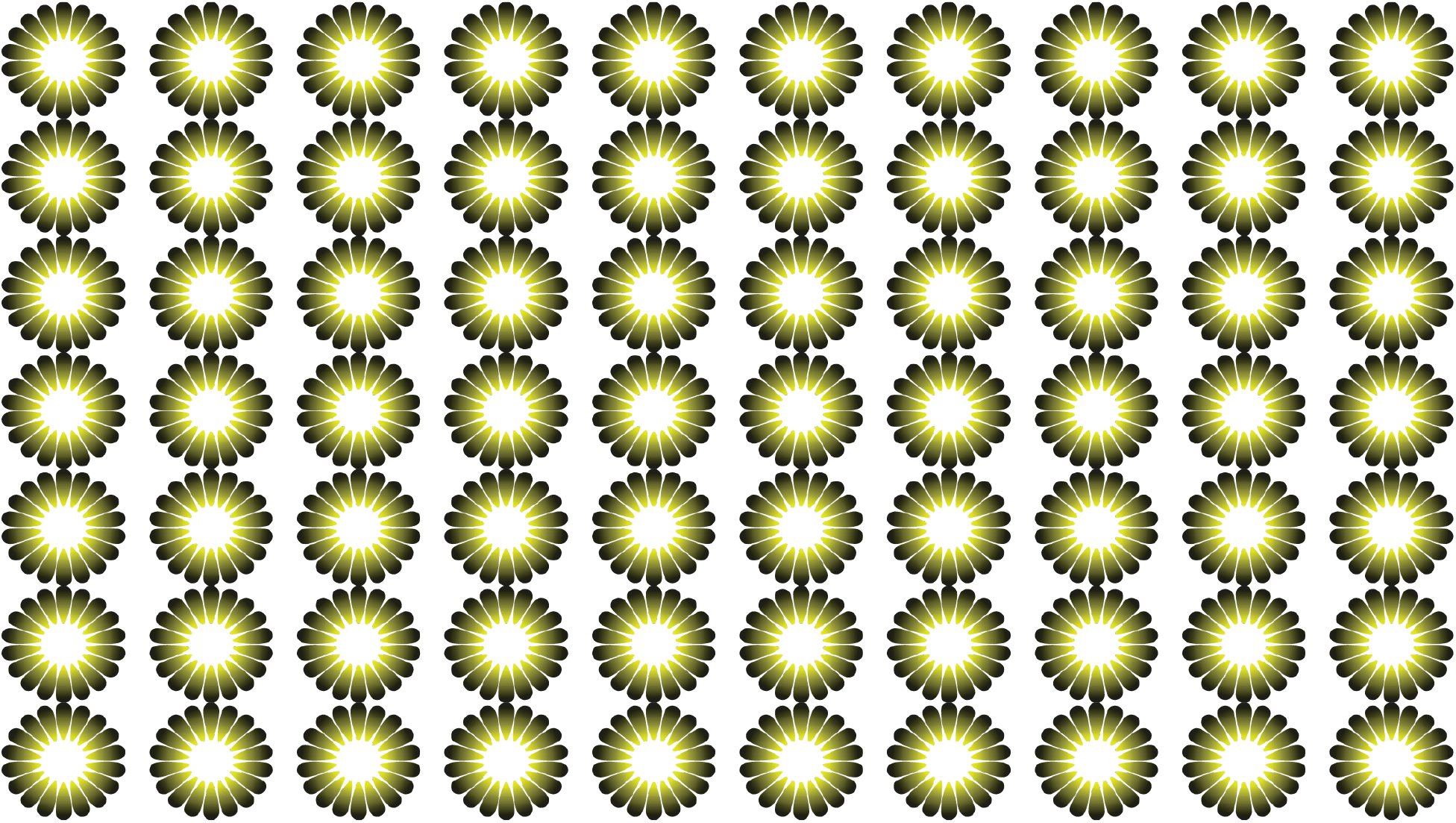
The yellow version of the 10° Asahi stimulus.

**Supplementary Figure 3.**
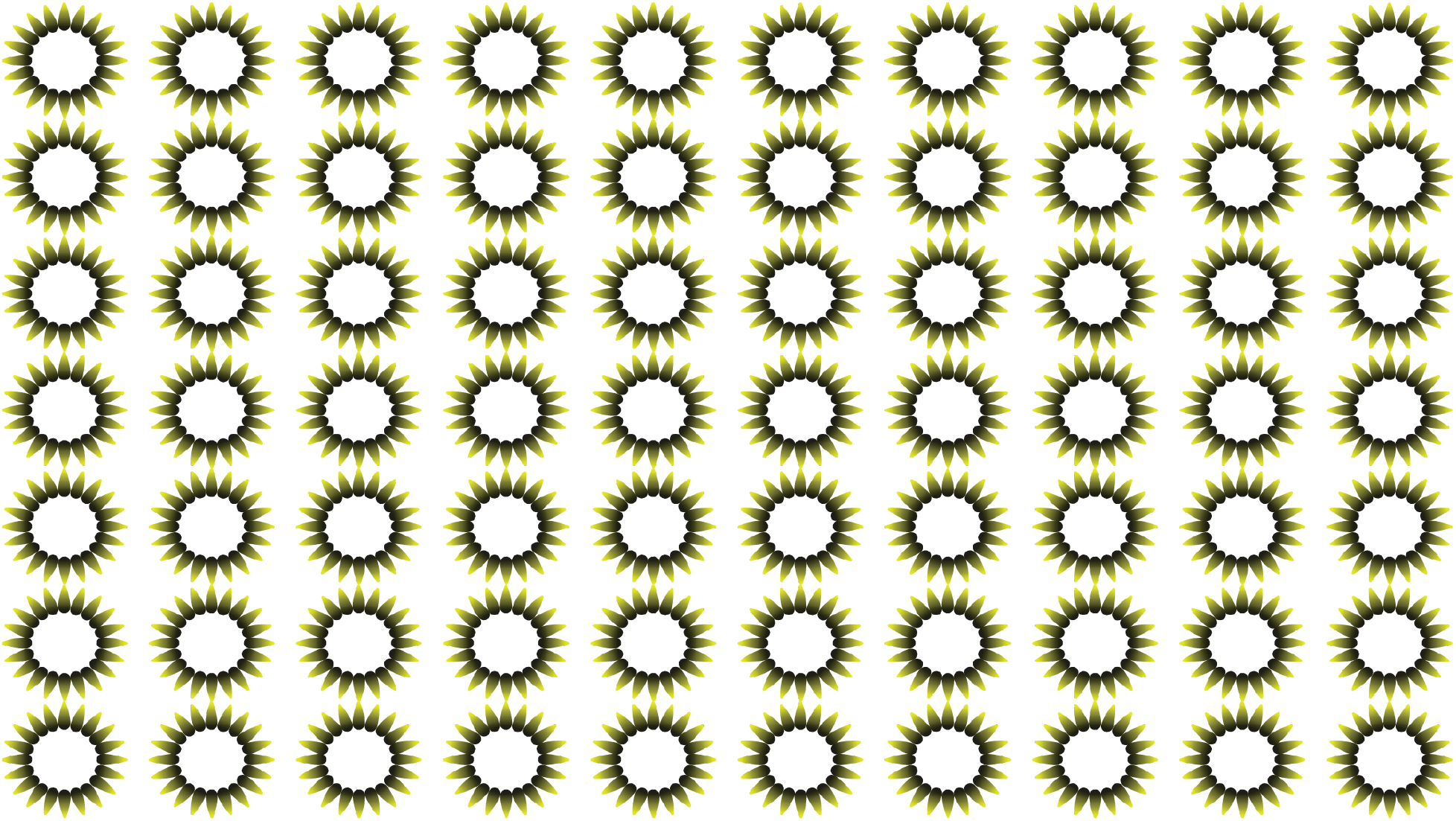
The yellow version of the 10° control stimulus.

**Supplementary Figure 4.**
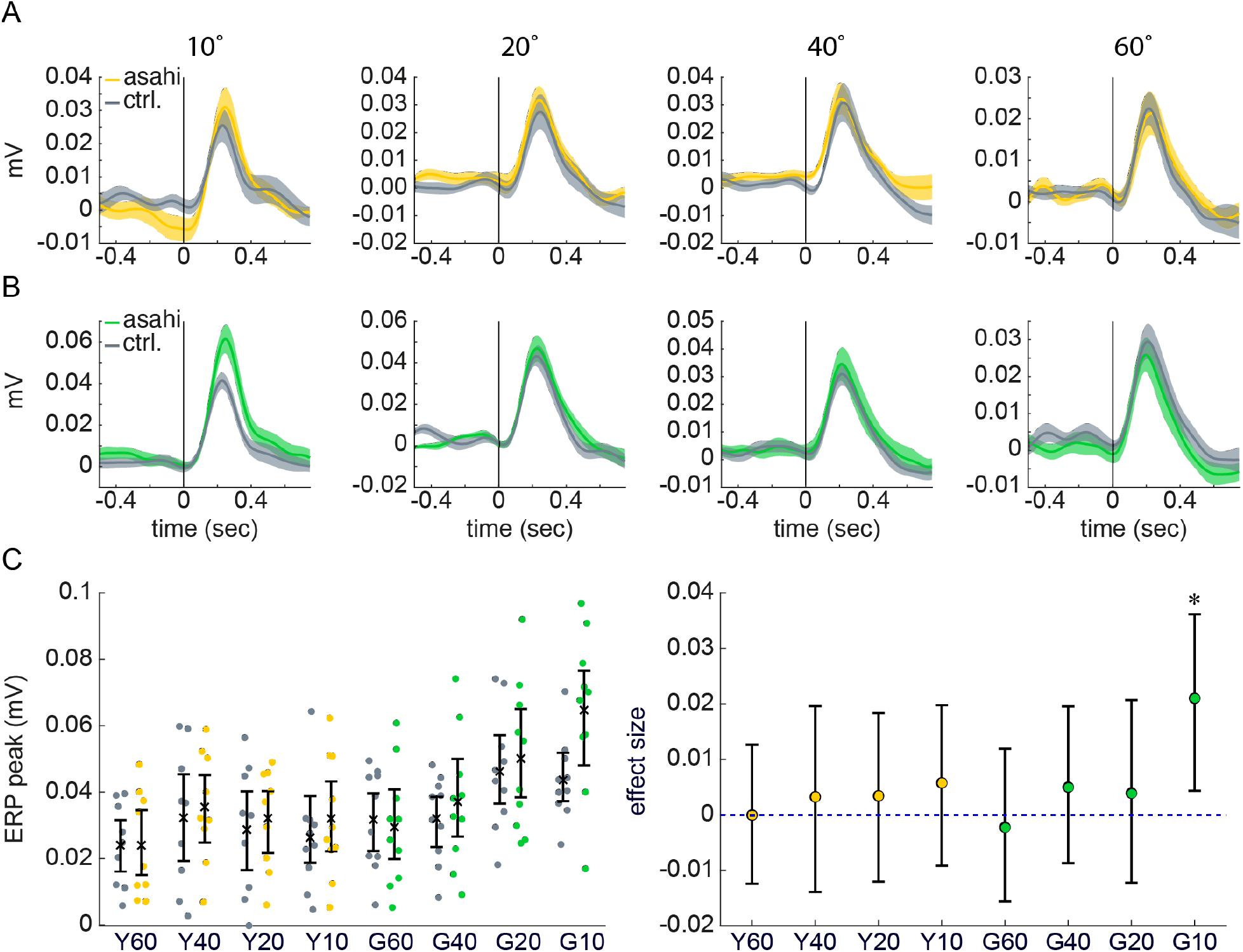
The ERP after each stimulus. **(A, B)** The SEM of the visual cortex EEG is plotted from 750 msec before and until 750 msec after stimulus onset. The grey line is the ERP around the control stimulus and the colored line is relative to the Asahi stimulus. Panel A shows the ERP to yellow stimuli and panel B shows the ERP to green stimuli. **(C)** The maximal potential in the ERP is plotted for each stimulus. Dots are individual rats. Note that one of the ten rats was not recorded in the yellow stimuli condition. The right panel shows the effect size between the Asahi stimulus and the control stimulus. The asterisk indicates support for the alternative hypothesis (BF10 > 3).

## Data sharing plan

All data and code will be shared in a Gnode repository upon publication.

## Acknowledgements

We thank Claudius Kratochwil for comments on the manuscript. We wish to thank the Finnish Grid and Cloud Infrastructure (FGCI) for supporting this project with computational and data storage resources. This work was funded by the Max Planck Society and the University of Helsinki (Helsinki Institute of Life Science).

## Author contributions

Conceptualization – NT; Data acquisition and curation – DV, IR; Formal analysis – DV, NT; Methodology – DV, NT; Project administration – NT; Supervision – NT; Visualization – NT; Writing – NT.

## Materials and Methods

### Subjects

Male, Lister-Hooded rats (140 grams to 190 grams) were obtained from Charles River. After a 7-day acclimation period, rats were implanted with a chamber and head-post and, in some cases, an EEG array. After implantation, rats were single housed. Experiments were carried out during the rats’ active phase (housing illumination from 7PM to 7AM). All procedures were carried out with the prior approval of local authorities and in compliance with the European Community Guidelines for the Care and Use of Laboratory Animals.

### Surgical procedures

The surgical procedure was identical to prior work^31^. Briefly, the rat was anesthetized using isoflurane and head-fixed using ear bars. We administered buprenorphine (0.06 mg/kg, s.c.), meloxicam (2.0 mg/kg, s.c.), enrofloxacin (10.0 mg/kg, s.c.), and lidocaine (0.5%, s.c. over skull) and waited 10 – 15 min (and for lack of response to paw pinch) before beginning surgical procedures. An EEG array (Neuronexus, CM32) was laid onto the cleaned and dried skull and fixed in place using dental cement (2-stage, powder/liquid Paladur). A custom-made skull implant was used for head-fixation (machine shop, Max Planck Institute for Biological Cybernetics). The implant was fixed onto the skull using UV-curing primer and dental cement (Tetric Evoflow, Dental Bauer). A craniotomy was made on the left occipital bone for a ground wire (99.9% pure silver). One end was flattened using an industrial press into a ~1-2 mm wide rectangle, which then was twisted into a roll to fit the craniotomy and inserted into the space between bone and dura. A rolled shape was used to increase the potential surface area in contact with CSF. The craniotomy was filled with viscous agar, which stabilized the wire and provided a conductive medium between the ground wire and the CSF. The other end of the ground wire was soldered to the ground wire of the electrode interface board of the EEG array. The wires and array were buried under dental cement (Paladur). The skin around the implant was glued to the implant using tissue glue (Histoacryl, B. Braun). Post surgical recovery lasted five days. During the first three days (surgery itself was counted as day one), the rat was injected either every 12 hours with buprenorphine or every 24 hours with meloxicam (same dosages as pre-operative). During the first five days, enrofloxacin was injected every 24 hours (same dosage as pre-operative). A rehydrating, nutritious, easily consumed, and palatable food was provided during recovery (DietGel Recovery, Clear H2O).

### Handling and habituation

Rats were handled daily for at least five minutes per day from the day of arrival in housing until the day of surgery, which was 7 days. Animals were neither food nor water restricted. Habituation consisted of one session (~25 minutes) of head-fixation on a freely-rotating treadmill in front of a computer screen. Rats were free to run or remain immobile and a mixture of locomotor activity was observed.

### Visual stimuli and stimulus presentation

Stimuli consisted of the Asahi stimulus, the control stimulus, and a grey screen used during the inter-stimulus interval. All stimuli were equiluminant (15 – 16 lux measured at the head-post). Stimuli were presented 50 cm from the rat’s head. The resolution was 1280 pixels by 720 pixels. Stimuli were created in Adobe Illustrator with a canvas set to match the screen resolution. These files were exported to JPEG at 72 ppi. The stimuli were presented using Psychtoolbox implemented in MATLAB. The stimuli were presented at four visual angles (10°, 20°, 40°, and 60°) and in two colors (yellow, 580 nm, HEX: #ffff00 and green, 508 nm, HEX: #00ff28). As many stimuli were placed on the screen as possible, thus covering the entire visual field or most of it. Stimuli were presented for a duration of 4 sec. The inter-stimulus intervals were drawn from a distribution with a flat hazard rate that spanned 4 to 8 sec with a 0.5 sec resolution.

### Pupillometry data acquisition and processing

Videos were recorded at 45 frames per second from the rat’s right eye with near-infrared illumination (Thor Labs LED, M850L3 and Thor Labs Collimation optics, COP4-B). Frames were acquired using a near-infrared camera (Allied Vision, G-046B) and variable zoom lens, fixed 3.3x zoom lens, and 0.25x zoom lens attachment (Polytec, 1-60135, 1-62831, 6044). Acquisition occurred over a GigE connection (MATLAB image processing toolbox). The camera provided a TTL pulse with each video frame. These TTL pulses were recorded directly into the neurophysiology system (Neuralynx).

We used an in-house custom algorithm and computer code to extract pupil size from the recorded video frames. The procedure is reviewed in detail in prior work^31^. Briefly, images were Gaussian blurred, converted into a binary image, and then subjected to edge detection, closed contour detection, and fitting of ellipses to the closed contours. In cases where the algorithm was not able to find an ellipse of an area bigger than predefined minimal allowed area (or smaller than maximal, respectively), then the value of the pupil in this frame was left blank. This was also applied to the frames which captured the animal blinking. Blank frames were linearly interpolated. The pupil detection algorithm was implemented using the OpenCV package in Python 3.7.

The pupil size was normalized to a pre-stimulus baseline (1 sec duration) by calculating a z-score. The z-score was calculated on each trial by subtracting the baseline mean from each pupil size data point and then dividing this array by the standard deviation of the baseline data points. The latency for pupil constriction was calculated as in prior work^20^. We first smoothed the pupil size with an 11 point, 2^nd^ order Savitzky-Golay filter (sgolayfilt in MATLAB). The signal was differentiated to obtain velocity and the velocity was lowpass filtered at 6 Hz with a 2^nd^ order Butterworth filter (filtfilt in MATLAB). This signal was then differentiated to obtain acceleration and the latency to constrict was defined as the time point with the largest negative acceleration.

### EEG signal acquisition and analysis

EEG signals were recorded using a flexible polyimide array with 32 platinum electrodes (Neuronexus, H32). Signals were recorded against animal ground, pre-amplified at the rat’s head (Neuralynx, HS-36), and then amplified and digitized at 32 kHz (Neuralynx, Digital Lynx SX). The analysis focused on bilateral electrodes placed over frontal cortex (locations relative to Bregma: 1.5 mm anterior, ±1.2 mm lateral; 3.6 mm anterior, ±1.2 mm lateral) and visual cortex (locations relative to Bregma: 5.0 mm posterior, ±1.5 mm lateral; 5.0 mm posterior, ±3.0 mm lateral; 5.0 mm posterior, ±4.4 mm lateral; 7.0 mm posterior, ±1.5 mm lateral; 7.0 mm posterior, ±3.0 mm lateral; 7.0 mm posterior, ±4.4 mm lateral). EEG signals were first low pass filtered at 5 Hz and then downsampled to 320 Hz. The entire signal was mean subtracted. EEG topographical plots were produced using the MATLAB command, scatteredInterpolant with natural neighbor interpolation. The contour plots (contourf function in MATLAB) used 50 levels.

### Statistics

We used estimation statistics and report effect sizes and the confidence intervals for effect sizes (DABEST toolbox in MATLAB^32,33^). Bayesian statistics were used for assessing evidence (or lack thereof) for the null hypothesis and for the alternative hypothesis^34^. Bayesian statistics were calculated in JASP software.

## References

1. Glaser, R., and Kiecolt-Glaser, J.K. (2005). Stress-induced immune dysfunction: implications for health. Nat Rev Immunol 5, 243–251. 10.1038/nri1571.

2. Mawdsley, J.E., and Rampton, D.S. (2005). Psychological stress in IBD: new insights into pathogenic and therapeutic implications. Gut 54, 1481. 10.1136/gut.2005.064261.

3. Poller, W.C., Downey, J., Mooslechner, A.A., Khan, N., Li, L., Chan, C.T., McAlpine, C.S., Xu, C., Kahles, F., He, S., et al. (2022). Brain motor and fear circuits regulate leukocytes during acute stress. Nature 607, 578–584. 10.1038/s41586-022-04890-z.

4. Geuter, S., Koban, L., and Wager, T.D. (2016). The Cognitive Neuroscience of Placebo Effects: Concepts, Predictions, and Physiology. Annu Rev Neurosci 40, 1–22. 10.1146/annurev-neuro-072116-031132.

5. Laeng, B., and Endestad, T. (2012). Bright illusions reduce the eye’s pupil. Proc National Acad Sci 109, 2162–2167. 10.1073/pnas.1118298109.

6. McDougal, D.H., and Gamlin, P.D. (2017). Comprehensive Physiology. Compr Physiol 5, 439–473. 10.1002/cphy.c140014.

7. Zavagno, D. (1997). Some New Luminance-Gradient Effects. Perception 28, 835–838. 10.1068/p2633.

8. Peirson, S.N., Brown, L.A., Pothecary, C.A., Benson, L.A., and Fisk, A.S. (2018). Light and the laboratory mouse. J Neurosci Meth 300. 10.1016/j.jneumeth.2017.04.007.

9. Beatty, J. (1982). Phasic Not Tonic Pupillary Responses Vary With Auditory Vigilance Performance. Psychophysiology 19, 167–172. 10.1111/j.1469-8986.1982.tb02540.x.

10. Breton-Provencher, V., and Sur, M. (2019). Active control of arousal by a locus coeruleus GABAergic circuit. Nat Neurosci 22, 218–228. 10.1038/s41593-018-0305-z.

11. Murphy, P.R., Robertson, I.H., Balsters, J.H., and O’connell, R.G. (2011). Pupillometry and P3 index the locus coeruleus-noradrenergic arousal function in humans. Psychophysiology 48, 1532–1543. 10.1111/j.1469-8986.2011.01226.x.

12. Euler, T., and Wässle, H. (1995). Immunocytochemical identification of cone bipolar cells in the rat retina. J Comp Neurol 361, 461–478. 10.1002/cne.903610310.

13. Prusky, G.T., Harker, K.T., Douglas, R.M., and Whishaw, I.Q. (2002). Variation in visual acuity within pigmented, and between pigmented and albino rat strains. Behav Brain Res 136, 339–348. 10.1016/s0166-4328(02)00126-2.

14. Artal, P., Tejada, P.H. de, Tedó, C.M., and Green, D.G. (1998). Retinal image quality in the rodent eye. Visual neuroscience 15, 597 605.

15. Dum, R.P., Levinthal, D.J., and Strick, P.L. (2019). The mind–body problem: Circuits that link the cerebral cortex to the adrenal medulla. Proc National Acad Sci 116, 26321–26328. 10.1073/pnas.1902297116.

16. Charbonneau, J.A., Maister, L., Tsakiris, M., and Bliss-Moreau, E. (2022). Rhesus monkeys have an interoceptive sense of their beating hearts. Proc National Acad Sci 119, e2119868119. 10.1073/pnas.2119868119.

17. Young, M.J., and Lund, R.D. (1994). The anatomical substrates subserving the pupillary light reflex in rats: Origin of the consensual pupillary response. Neuroscience 62, 481–496. 10.1016/0306-4522(94)90381-6.

18. Clarke, R.J., and Ikeda, H. (1985). Luminance and darkness detectors in the olivary and posterior pretectal nuclei and their relationship to the pupillary light reflex in the rat. Exp Brain Res 57, 224–232. 10.1007/bf00236527.

19. Fotiou, D.F., Brozou, C.G., Tsiptsios, D.J., Fotiou, A., Kabitsi, A., Nakou, M., Giantselidis, C., and Goula, A. (2007). Effect of age on pupillary light reflex: evaluation of pupil mobility for clinical practice and research. Electromyo Clin Neur 47, 11–22.

20. Bergamin, O., and Kardon, R.H. (2003). Latency of the Pupil Light Reflex: Sample Rate, Stimulus Intensity, and Variation in Normal Subjects. Investigative Opthalmology Vis Sci 44, 1546. 10.1167/iovs.02-0468.

21. Ellis, C.J. (1981). The pupillary light reflex in normal subjects. Brit J Ophthalmol 65, 754. 10.1136/bjo.65.11.754.

22. Purves, D., Williams, S.M., Nundy, S., and Lotto, R.B. (2004). Perceiving the Intensity of Light. Psychol Rev 111, 142–158. 10.1037/0033-295x.111.1.142.

23. Pak, A., Ryu, E., Li, C., and Chubykin, A.A. (2019). Top-Down Feedback Controls the Cortical Representation of Illusory Contours in Mouse Primary Visual Cortex. J Neurosci 40, 648–660. 10.1523/jneurosci.1998-19.2019.

24. Rossi, A.F., Rittenhouse, C.D., and Paradiso, M.A. (1996). The Representation of Brightness in Primary Visual Cortex. Science 273, 1104–1107. 10.1126/science.273.5278.1104.

25. Roe, A.W., Lu, H.D., and Hung, C.P. (2005). Cortical processing of a brightness illusion. Proc National Acad Sci 102, 3869–3874. 10.1073/pnas.0500097102.

26. Saeedi, A., Wang, K., Nikpourian, G., Bartels, A., Totah, N.K., Logothetis, N.K., and Watanabe, M. (2022). Mouse primary visual cortex neurons respond to the illusory “darker than black” in neon color spreading. Biorxiv, 2022.07.24.501311. 10.1101/2022.07.24.501311.

27. Panagiotaropoulos, T.I., Deco, G., Kapoor, V., and Logothetis, N.K. (2012). Neuronal Discharges and Gamma Oscillations Explicitly Reflect Visual Consciousness in the Lateral Prefrontal Cortex. Neuron 74, 924–935. 10.1016/j.neuron.2012.04.013.

28. Totah, N.K., Logothetis, N.K., and Eschenko, O. (2021). Synchronous spiking associated with prefrontal high gamma oscillations evokes a 5 Hz-rhythmic modulation of spiking in locus coeruleus. J Neurophysiol. 10.1152/jn.00677.2020.

29. Joshi, S., Li, Y., Kalwani, R.M., and Gold, J.I. (2016). Relationships between Pupil Diameter and Neuronal Activity in the Locus Coeruleus, Colliculi, and Cingulate Cortex. Neuron 89, 221–234. 10.1016/j.neuron.2015.11.028.

30. Luppi, P.-H., Aston-Jones, G., Akaoka, H., Chouvet, G., and Jouvet, M. (1995). Afferent projections to the rat locus coeruleus demonstrated by retrograde and anterograde tracing with cholera-toxin B subunit and Phaseolus vulgaris leucoagglutinin. Neuroscience 65, 119–160. 10.1016/0306-4522(94)00481-j.

31. Vasilev, D., Watanabe, M., Logothetis, N.K., and Totah, N.K. (2022). Focusing perceptual attention in one modality constrains subsequent learning in another modality. Biorxiv, 2022.01.22.477334. 10.1101/2022.01.22.477334.

32. Ho, J., Tumkaya, T., Aryal, S., Choi, H., and Claridge-Chang, A. (2019). Moving beyond P values: data analysis with estimation graphics. Nat Methods 16, 565–566. 10.1038/s41592-019-0470-3.

33. Calin-Jageman, R.J., and Cumming, G. (2019). Estimation for Better Inference in Neuroscience. Eneuro 6, ENEURO.0205-19.2019. 10.1523/eneuro.0205-19.2019.

34. Keysers, C., Gazzola, V., and Wagenmakers, E.-J. (2020). Using Bayes factor hypothesis testing in neuroscience to establish evidence of absence. Nat Neurosci 23, 788–799. 10.1038/s41593-020-0660-4.

